# Synchronization and metachronal waves of elastic cilia caused by transient viscous flow

**DOI:** 10.1101/2024.06.15.599160

**Authors:** Albert von Kenne, Sonja Schmelter, Holger Stark, Markus Bär

**Affiliations:** Physikalisch-Technische Bundesanstalt, Berlin 10587, Germany; Technische Universität Berlin, Berlin 10623 Germany

## Abstract

Hydrodynamic coordination of cilia is ubiquitous in biology. It is commonly modeled using the steady Stokes equations. The flow around ciliated cells, however, exhibits finite time vorticity diffusion, requiring a dynamical description. We present a model of elastic cilia coupled by transient viscous flow in the bulk fluid. Therein, vorticity diffusion impacts cilia coordination qualitatively and quantitatively. In particular, pairs of cilia synchronize in antiphase for long diffusion times. Moreover, metachronal waves occur in cilia chains larger than the viscous penetration depth, whereas global synchronization occurs in Stokes flow.

---

In eukaryotic life forms, hair-like cell extensions called cilia or flagella fulfill a wide variety of biological functions [1–4], ranging from swimming of microorganisms [5–7] to physiological flows in animals [8–10] and feeding of marine life [11, 12]. Their cyclic motions are often strikingly coordinated, showing perfectly synchronized beating [13] on small scales and traveling waves in the beating phase, called metachronal waves, on large scales [14–16]. Inspired by the ubiquity of cilia in nature, there has been considerable effort in micro-engineering to replicate biological principles in artificial cilia [17–19]. The emergence of coordinated motions is often attributed to hydrodynamic interactions [20–22], described by low Reynolds number fluid dynamics [23]. In the limit of zero Reynolds number, the viscous flow is governed by the Stokes equations [24], whose fundamental solutions can be readily used to model hydrodynamic coordination [25–27]. Indeed, measurements of average flow fields around flagellated cells can be fitted by Stokes flow [28]. Thus, most models of cilia coordination use this approximation to describe hydrodynamic interactions between cilia [29–65]. For coordinated motions to arise, the kinematic reversibility of Stokes flow must be broken [66], which is most notably achieved by the elasticity of cilia [32, 36, 67] or by specific variations in the forces driving their motion [37].

However, recent highly temporally resolved measurements with flagellated algae indicate that the flow surrounding microorganisms fundamentally differs from Stokes flow [68]. In particular, the oscillatory flow velocity measured at a distance from the algae is phase-shifted with respect to the flagella beating phase, and the spatial decay of the oscillatory flow velocity is faster than predicted for Stokes flow. Here, we address these observations through modeling a viscous flow by the time-dependent equations [69, 70]

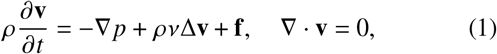

where **v** is the velocity field, *p* the pressure field, *ρ* the fluid density, *v* the kinematic viscosity, and **f** a body force density. An oscillatory forcing with period *T* causes an oscillatory flow that extends over the viscous penetration depth 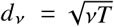 [71]. The transient flow spreads by vorticity diffusion, requiring the time *t*_*v*_ = *d*^2^ /*v* to diffuse a distance *d* from the forcing [69]. Using Eq. (1) to model the microscopic flow gives good agreement between theory and experiment [72]. When the acceleration in Eq. (1) is neglected, the Stokes equations follow. This term, however, breaks the kinematic reversibility of Stokes flow, which causes synchronization independent of an external symmetry breaking [73].

In this Letter, we study dynamics of interacting cilia beyond the limit of zero Reynolds number, taking into account the effect of vorticity diffusion. Our model consists of small spheres rotating elastically bound on a circular orbit in the bulk fluid [elastic rotors, Fig. 1(a)], coupled to each other by transient viscous flows caused by their motion. We show that vorticity diffusion qualitatively and quantitatively influences synchronization and metachronal waves. Contrary to the findings in Stokes flow [36], pairs of rotors with the same chirality switch to anti-phase synchronization [Fig. 1(b)] when the vorticity diffusion time and the rotation period are similar, i. e. when the reduced frequency *τ*_*v*_ = *d*^2^/(*vT*) is of order one. The finite diffusion time reveals itself as a phase-delayed coupling, as we will demonstrate by reducing the rotor dynamics to coupled phase oscillators. We will use the resulting phase-oscillator model to study metachronal waves in chains of rotors with periodic boundaries. In Stokes flow, metachronal waves with finite wavelengths are unstable [38, 45, 65], while in transient flow such waves become stable when the chain size is comparable to the penetration depth 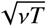. Depending on the orientation of the rotors in the chain, long wavelengths can be either stable or unstable.

**FIG. 1.**
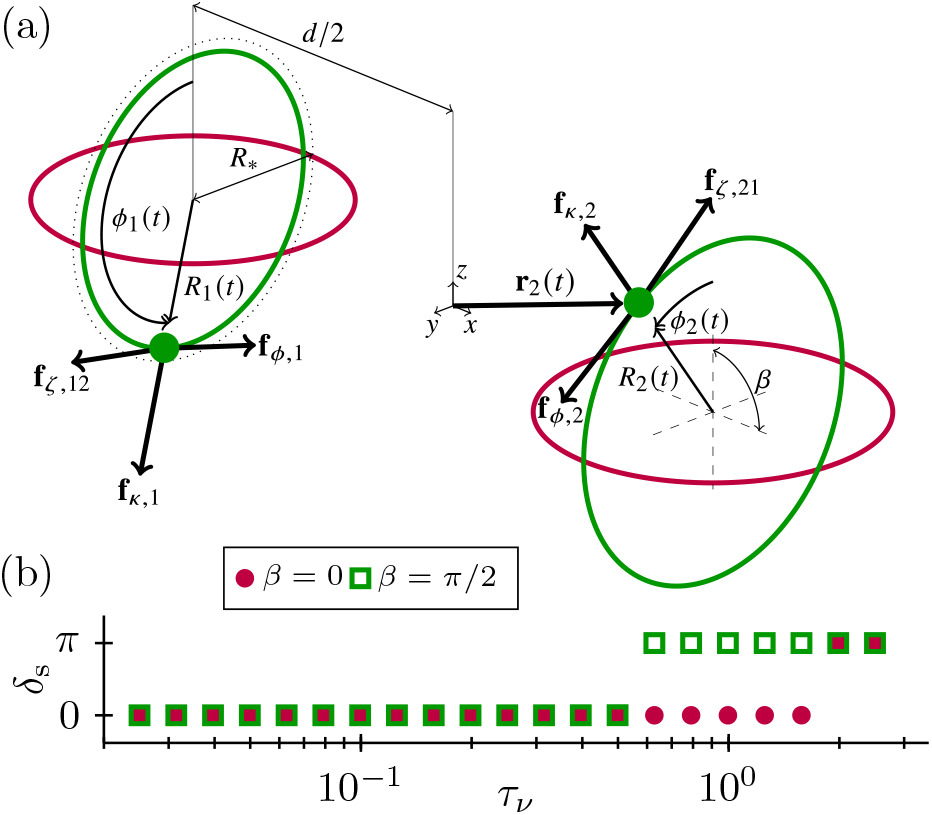
(a) Geometry of elastic rotors, whose orbits are either in the same plane (*β* = 0, purple), or in parallel planes (*β* = *π*/2, green). The forces exerted on the rotors and other symbols are explained in the main text. (b) Stationary phase difference δ_s_ = lim_*t*→∞_ *ϕ*_1_ (*t*) − *ϕ*_2_ (*t*), according to Eq. (3), as a function of the reduced frequency *τ*_*v*_ = *d*^2^ /(*vT*), which is the ratio of vorticity diffusion time and period of rotor motion.

We consider spherical rotors of radius *a*, as depicted in Fig. 1(a). Their positions are written as

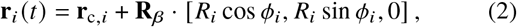

where *i* ∈ {1, 2}, **r**_c,*i*_ = (−1) ^*i*^ *d*/2**e**_*x*_, *d* is the center distance, **e**_*x*_ the unit vector in *x*-direction, *R*_*i*_ (*t*), *ϕ*_*i*_ (*t*) are radius and phase in polar coordinates, respectively, and the rotation matrix **R**_*β*_, see supplementary information (SI), orients the orbits either in the same plane (purple) or in parallel planes (green). The rotors are driven by a constant tangential force **f**_*ϕ,i*_ = *F***e**_*ϕ,i*_ of magnitude *F*, where **e**_*ϕ,i*_ = **R**_*β*_·[− sin *ϕ*_*i*_, cos *ϕ*_*i*_, 0]. A harmonic restoring force **f**_*k,i*_ = *k*(*R*_∗_ − *R*_*i*_)**e**_*R,i*_) with spring constant *κ*, reference radius *R*_∗_, and **e**_*R,i*_ = **R**_*β*_ · [cos *ϕ*_*i*_, sin *ϕ*_*i*_, 0 ], allows deviations from a pure circular motion. The fluid exerts a viscous drag force **f**_*ζ*, *ij*_ = *ζ*[**v** _*j*_ (*t*, **r**_*i*_) − d**r**_*i*_ /d*t*], where *ζ* = 6*πρva* and the background velocity field **v** _*j*_ is generated by the neighboring rotor. These forces put isolated rotors in circular motion with a period *T* = 2*πζ R*_∗_/*F*, while perturbations in radial direction relax on a timescale *t*_*k*_ = *ζ*/*κ*.

The dynamics of a small sphere in transient viscous flow is described by the Basset–Boussinesq–Oseen equation [74–77], which reduces to Stokes drag law if 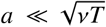 (SI). In this case, the rotors obey the force balance **f**_*ϕ,i*_ +**f**_*k,i*_ +**f**_*ζ*, *i j*_ = **0**, which gives the dynamics of the *i*th rotor as

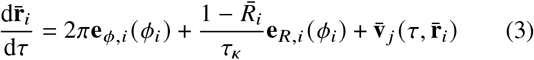

in terms of the dimensionless quantities *τ* = *t*/*T*, *τ*_*k*_ = *t*_*k*_ /*T*, 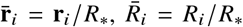 and 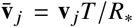. To describe the hydrodynamic interactions, we neglect the finite size of the orbits (*R*/*d* ≪ 1) and retain only the leading order in the interaction strength (*a*/*d* ≪ 1). Then, the velocity at the *i*th rotor, which is caused by the *j* th rotor, reads (SI)

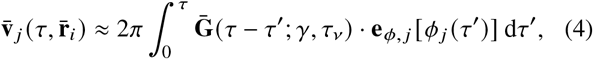

with 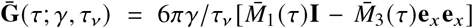, where *γ* = *a*/*d*, **I** is the identity matrix, and the memory kernels read 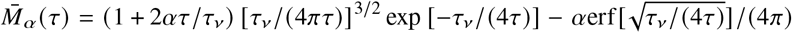 [73]. Thus, the rotor dynamics (3) is parametrized by the reduced frequencies *τ*_*k*_ and *τ*_*v*_, which indicate the softness of the rotor orbits and the vorticity diffusion time, respectively. The time-independent fundamental solution of the Stokes equations is called the Stokeslet. We call elastic rotors coupled by the Stokes equations the Stokeslet model.

We simulate the rotor dynamics (3) with the Euler method and calculate the velocity functional (4) in each time step with the trapezoidal rule (SI). Figure 1(b) shows that the rotors synchronize either inphase (δ_s_ = 0) or antiphase (δ_s_ = *π*), depending on *τ*_*v*_, while in the Stokeslet model (*τ*_*v*_→0) only in-phase synchronization occurs.

Figure 2(a) shows the diffusion of a vortex, emanated by an oscillating point force. The finite diffusion time causes a phase delay between the oscillatory flow velocity and the forcing [Fig. 2(b)]. We will see shortly that this phase delay causes the transition to anti-phase synchronization. The phase delay grows with distance [Fig. 2(c)] as the diffusion over a larger distance takes a longer time. The growth of the phase delay is larger perpendicular to the forcing, which motivates studying rotors either in the same or in parallel planes, respectively.

**FIG. 2.**
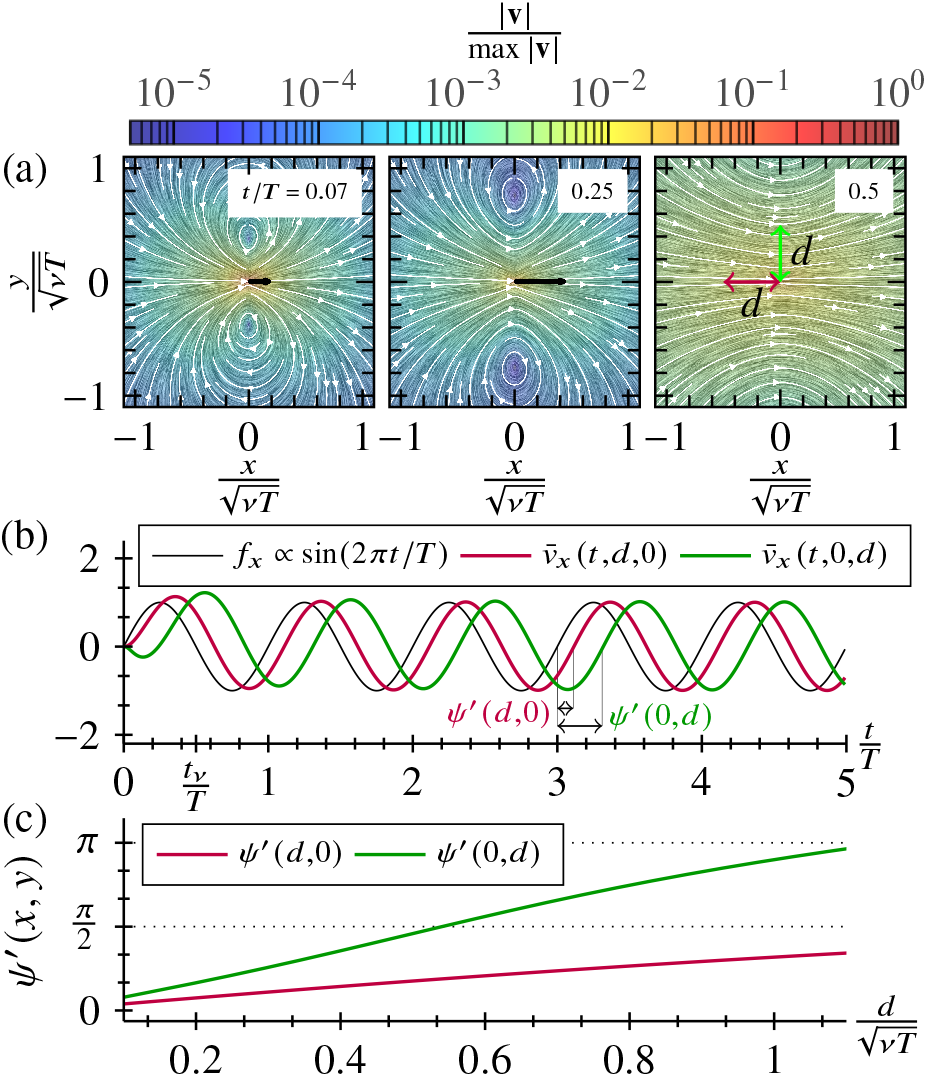
(a) Bulk fluid solution of Eq. (1) for a horizontal point force *f*_*x*_ ∝ sin (2*πt*/*T*), with period *T*. (b) Normalized velocity 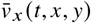 initiated by *f*_*x*_ at a distance *d* = (*vT*/2) ^1/2^, in vertical (green) and horizontal (purple) directions, corresponding to a vorticity diffusion time *t*_*v*_ = *T*/2. (c) Phase shift *ψ*^′^ between velocity and forcing in both directions.

The rotor dynamics (3) involve three inherent timescales: the period of rotor motion *T*, the elastic restoring time *t*_*κ*_, and the vorticity diffusion time *t*_*v*_. From the hydrodynamic coupling another timescale emerges: the synchronization time *t*_s_ =*O*(*d*/*a*). In the far field, the interaction is weak (*a*/*d* ≪ 1).

Thus, synchronization is slow (*t*_s_ ≫*T*, *t*_*k*_, *t*_*v*_). Hydrodynamic changes in the history of phases *ϕ*_*i*_ (*τ*) = *ϕ*_*i*_ (0) +2*πτ* + *O* (*a*/*d*) cause contributions of order *O*[( *a*/*d*)^2^] in the velocity (4). By neglecting these corrections and by assuming slow synchronization, we are able to derive an effective phase-oscillator model, which reflects the hydrodynamic coupling of rotors on the synchronization timescale (SI). For the phases of the two rotors we obtain the dynamics

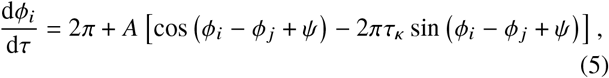

with amplitude 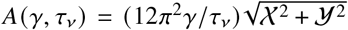 and phase delay *ψ* (*τ*_*v*_) = arctan (𝒴/𝒳). The effect of vorticity diffusion enters through the history integrals

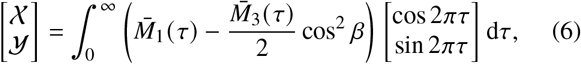

which, in general, we solve numerically (SI).

Figure 3(a) shows the memory kernel that weights the history of rotor motion [colored arrows in Fig. 3(b)] to generate the hydrodynamic coupling in the rotor model (3) and (4). The effective phase-oscillator model (5) replaces this history dependence by an effective force **f**_e,*j*_ ∝ **e**_*ϕ*, *j*_[*ϕ* _*j*_ (*τ*) − *ψ*] that acts on the fluid with a constant phase delay [black arrow in Fig. 3(b)]. This delay reflects the finite vorticity diffusion time. Furthermore, a reduced coupling amplitude relative to the Stokeslet model [compare straight and dotted lines in Fig. 3(b)] reflects cancellation of opposing directions in the history of rotor motion. On the timescale of synchronization, phase delay *ψ* and amplitude *A* can be calculated using the long time behavior of the coupling functionals (SI), which is represented by Eq. (6).

**FIG. 3.**
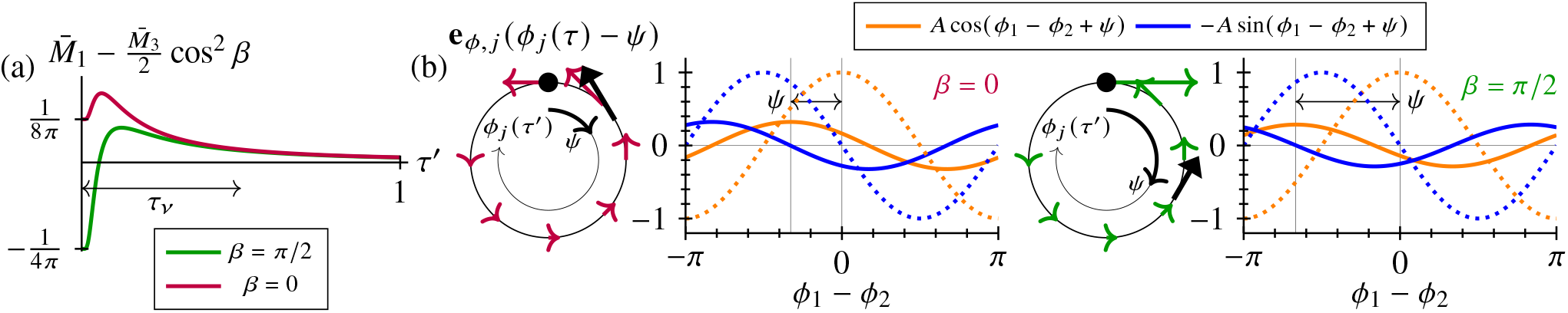
(a) Memory kernel of hydrodynamic coupling. (b) In the rotor model (3) and (4), the hydrodynamic coupling depends on the history of rotor motion weighted by the memory kernel (colored arrows). In the effective phase-oscillator model (5), the coupling is caused by an instantaneous force with phase delay *ψ* (black arrow). The resulting coupling functions (solid lines) are shifted and have a reduced amplitude compared to the Stokeslet model with *ψ* = 0 (dotted lines).

The collective phenomena of the rotor pair are encoded in the dynamics of the phase difference δ (*τ*) = *ϕ*_1_ (*τ*) − *ϕ*_2_ (*τ*) and the phase sum *σ*(*τ*) = *ϕ*_1_ (*τ*) + *ϕ*_2_ (*τ*). From the dynamics of phases (5) we derive

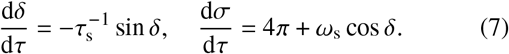

Here, 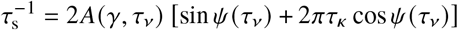 and *ω*_s_ = 2*A* (*γ*, *τ*_*v*_) [cos *ψ* (*τ*_*v*_) − 2*πτ*_*k*_ sin *ψ* (*τ*_*v*_) ], are the synchronization rate and the frequency gain, respectively. Note that the phase difference obeys the well-known Adler equation [78].

In the Stokeslet model, *ψ* = 0 so that 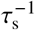 and *ω*_s_ are strictly positive, corresponding to in-phase synchronization and a co-operative increase in rotational frequency, respectively. The crucial difference in transient flow is that vorticity diffusion causes a finite phase delay *ψ* ≠ 0. Thus, 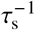 and *ω*_s_ can be negative, which causes anti-phase synchronization and a cooperative decrease in frequency, respectively. In particular, for soft rotors with 2*πτ*_*k*_ ≫ 1, the transition to anti-phase synchronization occurs at *ψ* ≈ *π*/2; for stiff rotors with 2*πτ*_*k*_ ≪ 1, it occurs at *ψ* ≈ *π*. For small phase delay *ψ* ≪ 1, we obtain 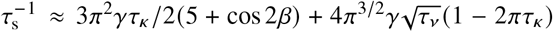. The leading order behavior of *ω*_s_ is given in the SI. Note that in the Stokeslet model (*τ*_*v*_ → 0), no synchronization occurs for rigid rotors (*τ*_*κ*_ → 0), while a phase delayed coupling can synchronize rigid rotors with a rate 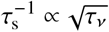 [73].

In general, vorticity diffusion delays the coupling of the rotors by the phase shown in Fig. 4(a), as a function of *τ*_*v*_. At the phase delays indicated by the dashed lines, 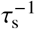 changes sign, as implied by the divergences in Fig. 4(c). As a result, the synchronization mode changes from inphase to antiphase and vice versa. The Stokeslet model predicts that the rotors synchronize in a time inversely proportional to *τ*_*k*_ [dotted lines in Fig. 4(c)]. This is counteracted by vorticity diffusion, which reduces the coupling amplitude at high frequency [Fig. 4(b)], leading to optimal synchronization at a finite frequency, either inphase (*β* = 0) or antiphase (*β* = *π*/2).

**FIG. 4.**
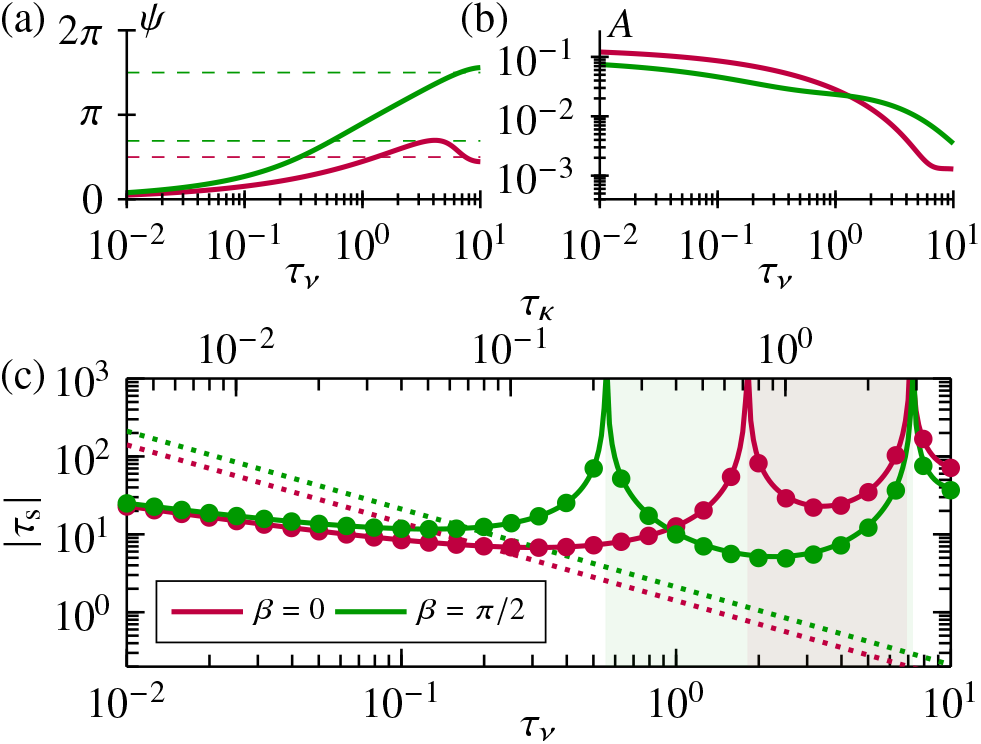
(a) Phase delay, (b) coupling amplitude, and (c) synchronization time as functions of reduced frequency *τ*_*v*_ and *τ*_*k*_. Purple and green solid lines and symbols denote data of the phase-oscillator model (5) and of the rotor model (3) and (4), respectively. The dashed lines in (a) indicate the delays corresponding to the divergences in (c). The colored regions in (c) correspond to antiphase synchronization. Dotted lines in (c) indicate the Stokeslet model, which always favours in-phase synchronization. Data are generated with *τ*_*v*_ /*τ*_*k*_ = 2.5, and *γ* = 1/50.

Next, we use the phase-oscillator model (5) to investigate metachronal waves in a chain of *N* rotors, with lattice constant *𝓁* (Fig. 5). The pair interactions in the chain are calculated by solving the history integrals [Eq. (6)] with *d*_*i j*_ = *𝓁* |*i*− *j*|, where *i, j* ∈ { 0, …, *N*−1} and *j* ≠ *i*. This defines the coupling amplitudes *A*_*ij*_ = *A* [*a*/*d*_*ij*_, (*d*_*ij*_)^2^ /(*vT*) ]. and phase delays *ψ*_*i j*_ = *ψ* [(*d*_*ij*_) ^2^ / (*vT*)]. From Eq. (5) we obtain the dynamic equations of the phase angles

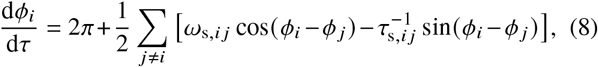

where 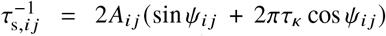 and *ω*_s,*i j*_ = 2*A*_*i j*_ (cos *ψ*_*i j*_ − 2*πτ*_*k*_ sin *ψ*_*ij*_). We impose periodic boundary conditions and a cutoff radius of half the system size.

**FIG. 5.**
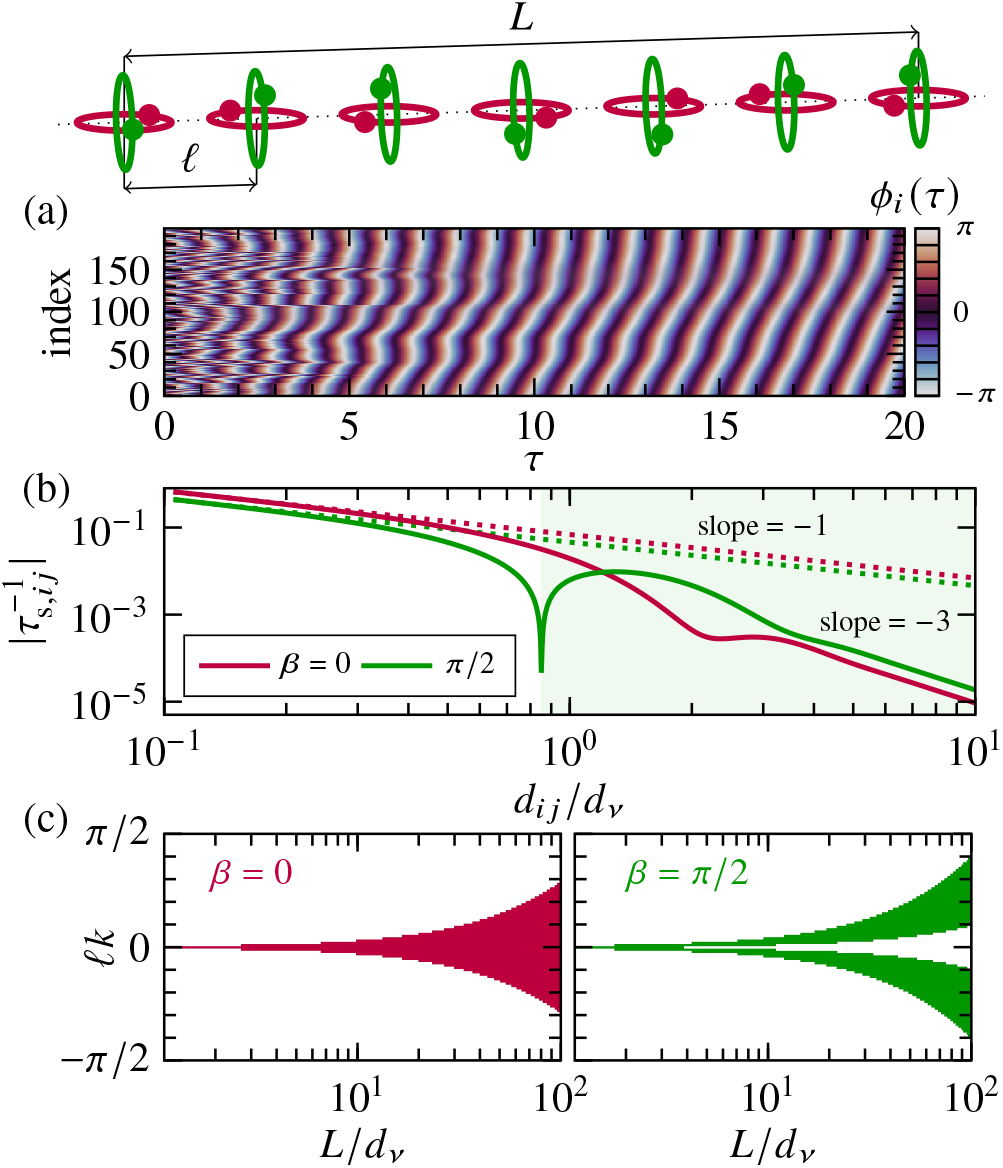
(a) Simulation of the phase dynamics (8) by the classical Runge-Kutta method with time step Δ*τ* = 0.01, random initial conditions, *β* = *π*/2, *a*/*𝓁* = 1/20, and *L*/*d*_*v*_ = 10. (b) Synchronization rate as a function of distance. The colored region indicates 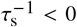 for *β* = *π* /2 and the dotted lines show data of the Stokeslet model. (c) Range of wave numbers *k* for which metachronal waves are stable (colored regions) as a function of *L*/*d*_*v*_. The data are generated with *N* = 200 and *τ*_*k*_ = 0.2.

In our model, metachronal waves are frequency-locked solutions of Eq. (8). The metachronal waves with wave number *k* are generated by constant phase shifts *ϕ*_*i*_ − *ϕ*_*i*+1_ = *𝓁k* with |*𝓁k* | = 2*πK*/*N*, where *K* is an integer. We perform a linear stability analysis of such a wave (SI) and obtain the growth rates of perturbations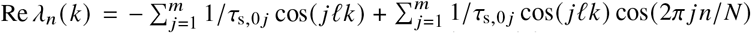, where *n* ∈ {0, …, *N* −1}, *m* = (*N* − 1) /2 for odd *N*, and *m* = *N* /2 − 1 for even *N*. A wave with wave number *k* is linearly stable if Re *λ*_*n*_ < 0 for all *n*. We discard Re *λ*_0_, which reflects the symmetry of Eq. (8) with respect to a global phase shift.

In the Stokeslet model, the synchronization rate decays slowly as 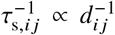 and metachronal waves are never stable [38, 45]. In our model, however, stable waves emerge from random initial conditions [Fig. 5(a)], because vorticity diffusion crucially affects the spatial decay of synchronization rate [Fig. 5(b)]. Beyond the penetration depth 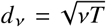, the decay crosses over to 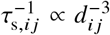, which stabilizes metachronal waves. In Stokes flow, a similar effect occurs at surfaces [38, 45, 65]. Furthermore, in our model the synchronization rate can become negative, as indicated by the divergence for *β* = *π*/ 2 in Fig. 5(b). Here, rotor pairs separated by a distance *d*_*ij*_ ⪆ *d*_*v*_ synchronize in antiphase. This renders the globally synchronized collective state unstable. Thus, the infinite wave-lenghts *𝓁k* = 0, which are exclusively stable in the Stokeslet model, are unstable [*β* = *π*/ 2 in Fig. 5(c)].

Whether vorticity diffusion causes metachronal waves depends on the ratio of chain length *L* = *N𝓁* and penetration depth *d*_*v*_, as seen in Fig. 5(c). Here, bands of stable wave numbers [colored regions in Fig. 5(c)] emerge from *𝓁k* = 0 as *L*/*d*_*v*_ increases. Notably, long wavelengths (small *𝓁k*) can either be stable (*β* = 0) or unstable (*β* = *π*/2).

In living matter, cilia are found on length scales ranging over several orders of magnitude, while the beating frequency is typically of the order of 10 Hz [1]. Unicellular organisms, such as *Paramecium*, have densely ciliated surfaces, extending over some hundred micrometers [5]. Large ciliated tissues, such as in brain ventricles [9], extends up to the centimetre scale. In many tissues, dense bundles of cilia are sparsely distributed throughout [79]. Whether finite-time vorticity diffusion qualitatively impacts cilia coordination can be estimated as follows: Antiphase synchronization requires at least a phase delay *ψ* (*τ*_*v*_) ≈ *π* / 2, which corresponds to a reduced frequency *τ*_*v*_ = *d*^2^ *f*/*v* ≈ 0.3 [Fig. 4(a)]. Thus, in water at body temperature with *v* ≈ 0.7 mm^2^ / s, rotors with a frequency *f*≈ 50 Hz would synchronize in antiphase when separated by a distance *d* ≈ 65*μ*m. In tissue, the typical separation between cilia bundles can be comparable [79]. Furthermore, the stability of metachronal waves is considerably affected by vorticity diffusion when *L*/*d*_*v*_ ⪆ 10 [Fig. 5(c)]. With *f* and *v* as before, the penetration depth is 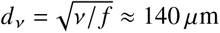. In this case, vorticity diffusion becomes relevant in systems of size *L* ⪆ 1.4 mm. However, in ciliated systems smaller than the penetration depth with cilia separations of only a few micrometer, no qualitative impact on the collective phenomena is expected. Nevertheless, vorticity diffusion has an effect. Namely, the synchronization rate 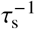 contains an *O*(*τ*_κ_) term reflecting orbit compliance, as well as an 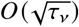 term reflecting vorticity diffusion. For nodal cilia, the elastic relaxation time *t*_*k*_ = *ζ*/*κ* is estimated between 2× 10^−4^ s and 2×10^−5^ s [36]. Using our analytical expression for 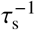 with *f* = 10 Hz, *d* = 10 *μ*m, *β* = 0, and *v* as before, we get a five to fifty fold increase in synchronization speed compared to the Stokeslet model.

Altogether we demonstrated that finite time vorticity diffusion impacts qualitatively and quantitatively the hydrodynamic coordination of elastic model cilia. The effect of the transient flow on synchronization and wave dynamics can be captured by a simple phase-oscillator model. This provides a convenient platform for more realistic modeling of hydrodynamic coordination in biological and artificial cilia. The theory presented can be tested experimentally by driving colloids on circular orbits with optical tweezers [27, 80, 81]. Modulating the driving forces throughout the orbits could reveal the interplay of vorticity diffusion and hydrodynamic coordination via variable forcing.

## References

[1] W. Gilpin, M. S. Bull, and M. Prakash, Nature Reviews Physics 2, 74 (2020).

[2] K. Y. Wan and G. Jékely, Philosophical Transactions of the Royal Society B 375, 20190148 (2020).

[3] M. L. Byron, D. W. Murphy, K. Katija, A. P. Hoover, J. Daniels, K. Garayev, D. Takagi, E. Kanso, B. J. Gemmell, M. Ruszczyk, et al., Integrative and Comparative Biology 61, 1674 (2021).

[4] K. Y. Wan and R. N. Poon, Current Opinion in Cell Biology 86, 102286 (2024).

[5] S. L. Tamm, The Journal of Cell Biology 55, 250 (1972).

[6] M. Polin, I. Tuval, K. Drescher, J. P. Gollub, and R. E. Goldstein, Science 325, 487 (2009).

[7] D. R. Brumley, M. Polin, T. J. Pedley, and R. E. Goldstein, Physical Review Letters 109, 28 (2012).

[8] S. Nonaka, H. Shiratori, Y. Saijoh, and H. Hamada, Nature 418, 96 (2002).

[9] R. Faubel, C. Westendorf, E. Bodenschatz, and G. Eichele, Science 353, 176 (2016).

[10] O. Mesdjian, C. Wang, S. Gsell, U. D’ortona, J. Favier, A. Viallat, and E. Loiseau, Physical Review Letters 129, 038101 (2022).

[11] W. Gilpin, V. N. Prakash, and M. Prakash, Nature Physics 13, 380 (2017).

[12] R. N. Poon, T. A. Westwood, H. Laeverenz-Schlogelhofer, E. Brodrick, J. Craggs, E. E. Keaveny, G. Jékely, and K. Y. Wan, Physical Review Research 5, L042037 (2023).

[13] K. Y. Wan and R. E. Goldstein, Proceedings of the National Academy of Sciences 113, E2784 (2016).

[14] E. W. Knight-Jones, The Quarterly Journal of Microscopical Science 95, 503 (1954).

[15] K. Y. Wan, Essays in Biochemistry 62, 829 (2018).

[16] C. Ringers, S. Bialonski, M. Ege, A. Solovev, J. N. Hansen, I. Jeong, B. M. Friedrich, and N. Jurisch-Yaksi, eLife 12, e77701 (2023).

[17] X. Dong, G. Z. Lum, W. Hu, R. Zhang, Z. Ren, P. R. Onck, and M. Sitti, Science Advances 6, eabc9323 (2020).

[18] V. Sahadevan, B. Panigrahi, and C.-Y. Chen, Micromachines 13, 735 (2022).

[19] T. ul Islam, Y. Wang, I. Aggarwal, Z. Cui, H. E. Amirabadi, H. Garg, R. Kooi, B. B. Venkataramanachar, T. Wang, S. Zhang, et al., Lab on a Chip 22, 1650 (2022).

[20] G. Taylor, Proceedings of the Royal Society A: Mathematical, Physical and Engineering Sciences 209, 447 (1951).

[21] J. Elgeti and G. Gompper, Proceedings of the National Academy of Sciences 110, 4470 (2013).

[22] D. R. Brumley, K. Y. Wan, M. Polin, and R. E. Goldstein, eLife 3, e02750 (2014), 1403.2100.

[23] E. Lauga, The fluid dynamics of cell motility, Vol. 62 (Cambridge University Press, 2020).

[24] E. M. Purcell, American Journal of Physics 45, 3 (1977).

[25] R. Golestanian, J. M. Yeomans, and N. Uchida, Soft Matter 7, 3074 (2011).

[26] J. Elgeti, R. G. Winkler, and G. Gompper, Reports on Progress in Physics 78, 056601 (2015).

[27] N. Bruot and P. Cicuta, Annual Review of Condensed Matter Physics 7, 323 (2016).

[28] K. Drescher, R. E. Goldstein, N. Michel, M. Polin, and I. Tuval, Physical Review Letters 105, 168101 (2010).

[29] S. Gueron, K. Levit-Gurevich, N. Liron, and J. J. Blum, Proceedings of the National Academy of Sciences 94, 6001 (1997).

[30] S. Gueron and K. Levit-Gurevich, Proceedings of the National Academy of Sciences 96, 12240 (1999).

[31] M. Cosentino Lagomarsino, B. Bassetti, and P. Jona, The European Physical Journal B-Condensed Matter and Complex Systems 26, 81 (2002).

[32] M. Reichert and H. Stark, European Physical Journal E 17, 493 (2005).

[33] A. Vilfan and F. Jülicher, Physical Review Letters 96, 058102 (2006).

[34] P. Lenz and A. Ryskin, Physical Biology 3, 285 (2006).

[35] B. Guirao and J.-F. Joanny, Biophysical Journal 92, 1900 (2007).

[36] T. Niedermayer, B. Eckhardt, and P. Lenz, Chaos 18, 1 (2008).

[37] N. Uchida and R. Golestanian, Physical Review Letters 106, 1 (2011).

[38] C. Wollin and H. Stark, The European Physical Journal E 34, 1 (2011).

[39] N. Osterman and A. Vilfan, Proceedings of the National Academy of Sciences 108, 15727 (2011).

[40] C. Mettot and E. Lauga, Physical Review E 84, 061905 (2011).

[41] M. Leoni and T. B. Liverpool, Physical Review E 85, 040901 (2012).

[42] J. Kotar, L. Debono, N. Bruot, S. Box, D. Phillips, S. Simpson, S. Hanna, and P. Cicuta, Physical Review Letters 111, 228103 (2013).

[43] A. Takamatsu, K. Shinohara, T. Ishikawa, and H. Hamada, Physical Review Letters 110, 248107 (2013).

[44] B. Nasouri and G. J. Elfring, Physical Review E 93, 033111 (2016).

[45] D. R. Brumley, N. Bruot, J. Kotar, R. E. Goldstein, P. Cicuta, and M. Polin, Physical Review Fluids 1, 1 (2016).

[46] R. E. Goldstein, E. Lauga, A. I. Pesci, and M. R. Proctor, Physical Review Fluids 1, 073201 (2016).

[47] A. Maestro, N. Bruot, J. Kotar, N. Uchida, R. Golestanian, and P. Cicuta, Communications Physics 1, 28 (2018).

[48] Y. Kawamura and R. Tsubaki, Physical Review E 97, 022212 (2018).

[49] K. Okumura, S. Nishikawa, T. Omori, T. Ishikawa, and A. Takamatsu, Physical Review E 97, 032411 (2018).

[50] N. Pellicciotta, E. Hamilton, J. Kotar, M. Faucourt, N. Delgehyr, N. Spassky, and P. Cicuta, Proceedings of the National Academy of Sciences 117, 8315 (2020).

[51] E. Hamilton, N. Pellicciotta, L. Feriani, and P. Cicuta, Philosophical Transactions of the Royal Society B 375, 20190152 (2020).

[52] F. O. Mannan, M. Jarvela, and K. Leiderman, Physical Review E 102, 033114 (2020).

[53] H. Guo, Y. Man, K. Y. Wan, and E. Kanso, Journal of The Royal Society Interface 18, 20200660 (2021).

[54] I. Tanasijević and E. Lauga, Physical Review E 103, 022403 (2021).

[55] F. Meng, R. R. Bennett, N. Uchida, and R. Golestanian, Proceedings of the National Academy of Sciences 118, 10.1073/pnas.2102828118 (2021).

[56] W. Liao and E. Lauga, Physical Review E 103, 042419 (2021).

[57] S. Maretvadakethope, Y. Hwang, and E. E. Keaveny, Physical Review Fluids 7, 053101 (2022).

[58] A. Solovev and B. M. Friedrich, New Journal of Physics 24, 013015 (2022).

[59] A. Solovev and B. M. Friedrich, Chaos: An Interdisciplinary Journal of Nonlinear Science 32 (2022).

[60] A. V. Kanale, F. Ling, H. Guo, S. Fürthauer, and E. Kanso, Proceedings of the National Academy of Sciences 119, e2214413119 (2022).

[61] M. Tatulea-Codrean and E. Lauga, Physical Review Letters 128, 208101 (2022).5

[62] B. Chakrabarti, S. Fürthauer, and M. J. Shelley, Proceedings of the National Academy of Sciences 119, e2113539119 (2022).

[63] D. J. Hickey, R. Golestanian, and A. Vilfan, Proceedings of the National Academy of Sciences 120, e2307279120 (2023).

[64] R. R. Bennett, arXiv preprint arXiv:2309.08274 (2023).

[65] A. von Kenne, M. Bär, and T. Niedermayer, Physical Review E 109, 054407 (2024).

[66] B. Friedrich, The European Physical Journal Special Topics 225, 2353 (2016).

[67] G. S. Klindt, C. Ruloff, C. Wagner, and B. M. Friedrich, Physical Review Letters 117, 1 (2016).

[68] D. Wei, P. G. Dehnavi, M.-E. Aubin-Tam, and D. Tam, Physical Review Letters 122, 124502 (2019).

[69] B. Eckhardt and J. Buehrle, The European Physical Journal Special Topics 157, 135 (2008).

[70] D. Wei, P. G. Dehnavi, M.-E. Aubin-Tam, and D. Tam, Journal of Fluid Mechanics 915, A70 (2021).

[71] L. D. Landau, E. M. Lifschitz, and W. Weller, Bd. 6, Hydrodynamik (1991).

[72] N. Bruot, P. Cicuta, H. Bloomfield-Gadêlha, R. E. Goldstein, J. Kotar, E. Lauga, and F. Nadal, Physical Review Fluids 6, 053102 (2021).

[73] M. Theers and R. G. Winkler, Physical Review E 88, 023012 (2013).

[74] A. B. Basset, A treatise on hydrodynamics: with numerous examples, Vol. 2 (Deighton, Bell and Company, 1888).

[75] J. Boussinesq, Théorie analytique de la chaleur: mise en harmonie avec la thermodynamique et avec la théorie mécanique de la lumière, Vol. 2 (Gauthier-Villars, 1903).

[76] C. W. Oseen, Hydrodynamik (Akademische Verlagsgesellschaft, 1927).

[77] M. R. Maxey and J. J. Riley, The Physics of Fluids 26, 883 (1983).

[78] R. Adler, Proceedings of the IRE 34, 351 (1946).

[79] F. Boselli, J. Jullien, E. Lauga, and R. E. Goldstein, Physical Review Letters 127, 198102 (2021).

[80] J. Kotar, M. Leoni, B. Bassetti, M. C. Lagomarsino, and P. Cicuta, Proceedings of the National Academy of Sciences 107, 7669 (2010).

[81] L. Damet, G. M. Cicuta, J. Kotar, M. C. Lagomarsino, and P. Cicuta, Soft Matter 8, 8672 (2012).

