## Supplemental Information for "Synchronization and metachronal waves of elastic cilia caused by transient viscous flow"

### I. GEOMETRY

The rotation matrix used in the main text to orient the rotors [Eq. (2)] by an angle $\beta$, is given by

$$\mathbf{R}_\beta = \begin{bmatrix} \cos\beta & 0 & \sin\beta \\ 0 & 1 & 0 \\ -\sin\beta & 0 & \cos\beta \end{bmatrix}. \tag{SI.1}$$

### II. HYDRODYNAMICS

#### A. Particle dynamics in transient viscous flow

The motion of a small sphere at position $\mathbf{r}(t)$ in transient viscous flow with background velocity $\mathbf{v}(t,\mathbf{r})$, under the absence of external forces (neglecting contributions of the order of Faxen corrections), is described by the Basset–Boussinesq–Oseen equation [1], which is written as

$$\frac{m_\mathrm{s}}{6\pi\rho\nu aT}\frac{\mathrm{d}^2\bar{\mathbf{r}}}{\mathrm{d}\tau^2} = \left(\bar{\mathbf{v}} - \frac{\mathrm{d}\bar{\mathbf{r}}}{\mathrm{d}\tau}\right) + \frac{V_\mathrm{s}}{12\pi\nu aT}\frac{\mathrm{d}}{\mathrm{d}\tau}\left(\bar{\mathbf{v}} - \frac{\mathrm{d}\bar{\mathbf{r}}}{\mathrm{d}\tau}\right) + \frac{a}{\sqrt{\pi\nu T}}\int_0^\tau (\tau-\tau')^{-1/2}\frac{\mathrm{d}}{\mathrm{d}\tau'}\left(\bar{\mathbf{v}} - \frac{\mathrm{d}\bar{\mathbf{r}}}{\mathrm{d}\tau'}\right)\mathrm{d}\tau', \tag{SI.2}$$

where $\tau = t/T$, $\bar{\mathbf{r}} = \mathbf{r}/R_*$, $\bar{\mathbf{v}} = \mathbf{v}T/R_*$, $\nu$ denotes kinematic viscosity, $\rho$ fluid density, $T$ period of motion, $R_*$ reference radius of circular motion, $m_\mathrm{s}$ is the mass of the sphere, $V_\mathrm{s}$ its volume, and $a$ its radius. By Eq. (SI.2), we see that the corrections to Stokes drag law can be neglected if $m_\mathrm{s} \ll 6\pi\rho\nu aT$, $V_\mathrm{s} \ll 12\pi a\nu T$, and $a \ll \sqrt{\pi\nu T}$. Assuming that the sphere density $\rho_\mathrm{s}$ is similar to the fluid density $\rho$, the essential constraint is $a \ll \sqrt{\nu T}$.

#### B. Fundamental solution of transient viscous flow

The bulk fluid solution of the equations of transient viscous flow [Eq. (1) in the main text] for a point force $\mathbf{f}\delta(\mathbf{r}-\mathbf{r_f})$, where $\mathbf{r_f}(t)$ is the position of the point force, is given by

$$\mathbf{v}(t,\mathbf{r}) = \int_0^t \mathbf{G}[t-t', \mathbf{r}-\mathbf{r_f}(t')]\cdot\mathbf{f}(t')\mathrm{d}t' \tag{SI.3}$$

with the Green's function [2]

$$\mathbf{G}(t,\mathbf{r}) = \frac{1}{\rho}\left[M_1(t,r)\mathbf{I} - M_3(t,r)\frac{\mathbf{rr}}{r^2}\right], \tag{SI.4}$$

where $r = |\mathbf{r}|$, $\mathbf{rr} = [r_i r_j]$, $\mathbf{I}$ is the identity matrix, and the hydrodynamic memory kernels $M_1$ and $M_3$ are defined as

$$M_\alpha(t,r) = \left[1 + \frac{2\alpha\nu t}{r^2}\right]\frac{1}{(4\pi\nu t)^{3/2}}\exp\left(-\frac{r^2}{4\nu t}\right) - \frac{\alpha}{4\pi r^3}\mathrm{erf}\left(\frac{r}{(4\nu t)^{1/2}}\right) \tag{SI.5}$$

with the error function $\mathrm{erf}(z) = 2/\sqrt{\pi}\int_0^z \exp(-t^2)\mathrm{d}t$.

For hydrodynamic interactions between the two rotors shown in Fig. 1(a) of the main text, the background velocity at the $i$th rotor is caused by the $j$th rotor. It can be calculated according to

$$\mathbf{v}_j(t,\mathbf{r}_i) = \int_0^t \mathbf{G}[t-t', \mathbf{r}_i(t)-\mathbf{r}_j(t')]\cdot[-\mathbf{f}_{\zeta,ji}(t')]\mathrm{d}t'. \tag{SI.6}$$

Here $i \in \{1,2\}$, $j \neq i$, $\mathbf{r}_i(t) = \mathbf{r}_{\mathrm{c},i} + \mathbf{R}_\beta\cdot[R_i(t)\cos\phi_i(t), R_i(t)\sin\phi_i(t), 0]$, $\mathbf{r}_{\mathrm{c},i} = (-1)^i d/2\mathbf{e}_x$, $d$ is the center distance, $\mathbf{e}_x$ the unit vector in $x$-direction, and $\mathbf{f}_{\zeta,ji} = \zeta[\mathbf{v}_i - \mathrm{d}\mathbf{r}_j/\mathrm{d}t]$, with $\zeta = 6\pi\rho\nu a$.

We assume the scaling separations $a/d \ll 1$, $R/d \ll 1$ and make two approximations: (i) We develop the Green's function [Eq. (SI.4)] in powers of $R/d$ and keep only the zeroth order. (ii) We only retain the leading order interaction terms in $a/d \ll 1$. Thus, to calculate the background velocity $\mathbf{v}_j$, we set $d\mathbf{r}_j/dt \approx 2\pi R_*/T\mathbf{e}_{\phi,j}$ (where $\mathbf{e}_{\phi,j} = \mathbf{R}_\beta \cdot [-\sin\phi_j, \cos\phi_j, 0]$) and neglect the hydrodynamic feedback in the viscous reaction force. Hence, we use $-\mathbf{f}_{\zeta,j} \approx 2\pi\zeta R_*/T\mathbf{e}_{\phi,j}$ in Eq. (SI.6), together with the Green's function to the zeroth order in $R/d \ll 1$, to obtain the dimensionless background velocity $\bar{\mathbf{v}}_j = \mathbf{v}_j T/R_*$ at the $i$th rotor as

$$\bar{\mathbf{v}}_j(\tau, \bar{\mathbf{r}}_i) \approx 2\pi \int_0^\tau \bar{\mathbf{G}}(\tau - \tau'; \gamma, \tau_\nu) \cdot \mathbf{e}_{\phi_j}[\phi_j(\tau')]\mathrm{d}\tau', \tag{SI.7}$$

where the normalized far field Green's function $\bar{\mathbf{G}} = T\zeta\mathbf{G}$ reads

$$\bar{\mathbf{G}}(\tau; \gamma, \tau_\nu) = \frac{6\pi\gamma}{\tau_\nu}\left[\bar{M}_1(\tau)\mathbf{I} - \bar{M}_3(\tau)\mathbf{e}_x\mathbf{e}_x\right] \tag{SI.8}$$

with $\gamma = a/d$ and $\tau_\nu = t_\nu/T$, where $t_\nu = d^2/\nu$ is the vorticity diffusion time. The normalized memory kernels are given by

$$\bar{M}_\alpha(\tau) = \left(1 + \frac{2\alpha\tau}{\tau_\nu}\right)\left(\frac{\tau_\nu}{4\pi\tau}\right)^{3/2}\exp\left(-\frac{\tau_\nu}{4\tau}\right) - \frac{\alpha}{4\pi}\mathrm{erf}\left(\sqrt{\frac{\tau_\nu}{4\tau}}\right). \tag{SI.9}$$

### III. PHASE OSCILLATOR DYNAMICS ON THE SYNCHRONIZATION TIMESCALE

#### A. Dynamic equations

In the tangential and radial direction, respectively, the rotor dynamics [Eq. (3) of the main text] read

$$\bar{R}_i(\tau)\frac{\mathrm{d}\phi_i(\tau)}{\mathrm{d}\tau} = 2\pi + \mathcal{T}_{ij}\left\{\phi_j(\tau)\right\}, \tag{SI.10}$$

$$\frac{\mathrm{d}\bar{R}_i(\tau)}{\mathrm{d}\tau} = \frac{1 - \bar{R}_i(\tau)}{\tau_\kappa} + \mathcal{R}_{ij}\left\{\phi_j(\tau)\right\} \tag{SI.11}$$

with $\bar{R}_i = R_i/R_*$ and $\tau_\kappa = t_\kappa/T$, where $t_\kappa$ is the elastic relaxation time. The tangential and radial coupling functionals $\mathcal{T}_{ij} = \mathbf{e}_{\phi,i} \cdot \bar{\mathbf{v}}_j(\tau, \bar{\mathbf{r}}_i)$ and $\mathcal{R}_{ij} = \mathbf{e}_{R,i} \cdot \bar{\mathbf{v}}_j(\tau, \bar{\mathbf{r}}_i)$ [where $\mathbf{e}_{R,i} = \mathbf{R}_\beta \cdot (\cos\phi_i, \sin\phi_i, 0)$], respectively, are given by

$$\mathcal{T}_{ij}\left\{\phi_j(\tau)\right\} = \frac{12\pi^2\gamma}{\tau_\nu}\int_0^\tau \left(\bar{M}_1(\tau - \tau') - \frac{\bar{M}_3(\tau - \tau')}{2}\cos^2\beta\right)\cos\left[\phi_i(\tau) - \phi_j(\tau')\right]\mathrm{d}\tau' + \frac{12\pi^2\gamma}{\tau_\nu}\int_0^\tau \frac{\bar{M}_3(\tau - \tau')}{2}\cos^2\beta\cos\left[\phi_i(\tau) + \phi_j(\tau')\right]\mathrm{d}\tau' \tag{SI.12}$$

and

$$\mathcal{R}_{ij}\left\{\phi_j(\tau)\right\} = \frac{12\pi^2\gamma}{\tau_\nu}\int_0^\tau \left(\bar{M}_1(\tau - \tau') - \frac{\bar{M}_3(\tau - \tau')}{2}\cos^2\beta\right)\sin\left[\phi_i(\tau) - \phi_j(\tau')\right]\mathrm{d}\tau' + \frac{12\pi^2\gamma}{\tau_\nu}\int_0^\tau \frac{\bar{M}_3(\tau - \tau')}{2}\cos^2\beta\sin\left[\phi_i(\tau) + \phi_j(\tau')\right]\mathrm{d}\tau'. \tag{SI.13}$$

We reduce the integro-differential equations [Eqs. (SI.10) and (SI.11)] to ordinary differential equations. For this, note the following: (i) The history of phases $\phi_i(\tau) = \phi(0) + 2\pi\tau + O(a/d)$ is changed from its intrinsic form by weak hydrodynamic interactions ($\gamma = a/d \ll 1$). The corrections to the history functionals [Eqs. (SI.12) and (SI.13)] induced by hydrodynamic changes in the history of phases are a higher order effect in $a/d$. (ii) In the far field, hydrodynamic synchronization is slow compared to the vorticity diffusion time $\tau_\mathrm{s} = t_\mathrm{s}/T \gg \tau_\nu$, where $t_\mathrm{s}$ is the synchronization time.

By using (i), we make the approximation $\phi_i(\tau) \approx \phi_i(0) + 2\pi\tau$ in the history integrals. Thus, we can use the relation between initial phase and instantaneous phase to write

$$\phi_i(\tau) - \phi_j(\tau') \approx \phi_i(\tau) - \phi_j(0) - 2\pi\tau' \iff \phi_i(\tau) - \phi_j(\tau) + 2\pi(\tau - \tau'), \tag{SI.14}$$

$$\phi_i(\tau) + \phi_j(\tau') \approx \phi_i(\tau) + \phi_j(0) + 2\pi\tau' \iff \phi_i(\tau) + \phi_j(\tau) - 2\pi(\tau - \tau'). \tag{SI.15}$$

Furthermore, we are only interested in the dynamics at the slow timescale of synchronization. By recalling (ii), which says that the synchronization timescale is a slow scale, we can take the limit $\tau \to \infty$ in the history functionals to extract the long-time coupling. With (i) and (ii) the functional (SI.12) reads

$$\lim_{\tau\to\infty} \mathcal{T}_{ij}\left\{\phi_j(0)+2\pi\tau\right\} = \frac{12\pi^2\gamma}{\tau_\nu} \lim_{\tau\to\infty} \int_0^\tau \left(\bar{M}_1(\tau-\tau') - \frac{\bar{M}_3(\tau-\tau')}{2}\cos^2\beta\right)\cos\left[\phi_i(\tau)-\phi_i(\tau)+2\pi(\tau-\tau')\right]\mathrm{d}\tau' + \frac{12\pi^2\gamma}{\tau_\nu} \lim_{\tau\to\infty} \int_0^\tau \frac{\bar{M}_3(\tau-\tau')}{2}\cos^2\beta\cos\left[\phi_i(\tau)+\phi_i(\tau)-2\pi(\tau-\tau')\right]\mathrm{d}\tau'. \quad \text{(SI.16)}$$

Using trigonometric identities, we can rewrite Eq. (SI.16) as

$$\lim_{\tau\to\infty} \mathcal{T}_{ij}\left\{\phi_j(0)+2\pi\tau\right\} = \frac{12\pi^2\gamma}{\tau_\nu}\left[\mathcal{X}\cos\delta_{ij}(\tau) - \mathcal{Y}\sin\delta_{ij}(\tau) + \bar{\mathcal{X}}\cos\sigma_{ij}(\tau) + \bar{\mathcal{Y}}\sin\sigma_{ij}(\tau)\right], \quad \text{(SI.17)}$$

where $\delta_{ij}(\tau) \equiv \phi_i(\tau)-\phi_j(\tau)$, $\sigma_{ij}(\tau) \equiv \phi_i(\tau)+\phi_j(\tau)$ and

$$\mathcal{X} \equiv \lim_{\tau\to\infty} \int_0^\tau \left(\bar{M}_1(\tau-\tau') - \frac{\bar{M}_3(\tau-\tau')}{2}\cos^2\beta\right)\cos\left[2\pi(\tau-\tau')\right]\mathrm{d}\tau', \quad \text{(SI.18)}$$

$$\mathcal{Y} \equiv \lim_{\tau\to\infty} \int_0^\tau \left(\bar{M}_1(\tau-\tau') - \frac{\bar{M}_3(\tau-\tau')}{2}\cos^2\beta\right)\sin\left[2\pi(\tau-\tau')\right]\mathrm{d}\tau', \quad \text{(SI.19)}$$

$$\bar{\mathcal{X}} \equiv \lim_{\tau\to\infty} \int_0^\tau \frac{\bar{M}_3(\tau-\tau')}{2}\cos^2\beta\cos\left[2\pi(\tau-\tau')\right]\mathrm{d}\tau', \quad \text{(SI.20)}$$

$$\bar{\mathcal{Y}} \equiv \lim_{\tau\to\infty} \int_0^\tau \frac{\bar{M}_3(\tau-\tau')}{2}\cos^2\beta\sin\left[2\pi(\tau-\tau')\right]\mathrm{d}\tau'. \quad \text{(SI.21)}$$

Thus, the dependence of the coupling functional on instantaneous phase and phase history, respectively, factorizes. Substituting $\xi = \tau - \tau'$ and taking the limit in Eqs. (SI.18) and (SI.19), yields

$$\begin{bmatrix}\mathcal{X}\\ \mathcal{Y}\end{bmatrix} = \int_0^\infty \left(\bar{M}_1(\xi) - \frac{\bar{M}_3(\xi)}{2}\cos^2\beta\right)\begin{bmatrix}\cos(2\pi\xi)\\ \sin(2\pi\xi)\end{bmatrix}\mathrm{d}\xi, \quad \text{(SI.22)}$$

which is Eq. (6) of the main text.

Using more trigonometric identities, we can write Eq. (SI.17) as

$$\lim_{\tau\to\infty} \mathcal{T}_{ij}\left\{\phi_j(0)+2\pi\tau\right\} = A\cos\left[\delta_{ij}(\tau)+\psi\right] + \bar{A}\cos\left[\sigma_{ij}(\tau)-\bar{\psi}\right], \quad \text{(SI.23)}$$

where the amplitudes $A$, $\bar{A}$ and phase delays $\psi$, $\bar{\psi}$ are given by

$$A(\gamma,\tau_\nu) = \frac{12\pi^2\gamma}{\tau_\nu}\sqrt{\mathcal{X}^2+\mathcal{Y}^2}, \quad \text{(SI.24)}$$

$$\bar{A}(\gamma,\tau_\nu) = \frac{12\pi^2\gamma}{\tau_\nu}\sqrt{\bar{\mathcal{X}}^2+\bar{\mathcal{Y}}^2}, \quad \text{(SI.25)}$$

$$\psi(\tau_\nu) = \arctan\left(\frac{\mathcal{Y}}{\mathcal{X}}\right), \quad \text{(SI.26)}$$

$$\bar{\psi}(\tau_\nu) = \arctan\left(\frac{\bar{\mathcal{Y}}}{\bar{\mathcal{X}}}\right). \quad \text{(SI.27)}$$

Using the same approximations as before and repeating the calculation for the history functional (SI.13), we obtain

$$\lim_{\tau\to\infty} \mathcal{R}_{ij}\left\{\phi_j(0)+2\pi\tau\right\} = A\sin\left[\delta_{ij}(\tau)+\psi\right] + \bar{A}\sin\left[\sigma_{ij}(\tau)-\bar{\psi}\right]. \quad \text{(SI.28)}$$

With Eqs. (SI.23) and (SI.28) we can write ordinary differential equations for the phase $\phi_i(\tau)$ and radius $\bar{R}_i(\tau)$, describing the long-time dynamics of the rotors as

$$\bar{R}_i(\tau)\frac{\mathrm{d}\phi_i(\tau)}{\mathrm{d}\tau} = 2\pi + A\cos\left[\delta_{ij}(\tau)+\psi\right] + \bar{A}\cos\left[\sigma_{ij}(\tau)-\bar{\psi}\right], \quad \text{(SI.29)}$$

$$\frac{\mathrm{d}\bar{R}_i(\tau)}{\mathrm{d}\tau} = \frac{1-\bar{R}_i}{\tau_\kappa} + A\sin\left[\delta_{ij}(\tau)+\psi\right] + \bar{A}\sin\left[\sigma_{ij}(\tau)-\bar{\psi}\right]. \quad \text{(SI.30)}$$

Eqs. (SI.29) and (SI.30), respectively, involve coupling contributions on a fast and a slow timescale. The phase difference changes on the timescale of synchronization $\mathrm{d}\delta_{ij}/\mathrm{d}\tau = O(\tau_s^{-1})$, but the phase sum changes on the period of motion $\mathrm{d}\sigma_{ij}/\mathrm{d}\tau = O(2\pi)$. We extract the dynamics on the timescale of synchronization by removing the fast coupling variations through cycle-averaging Eqs. (SI.29) and (SI.30). Therefore, we average over one period as $\langle\cdot\rangle = \int_0^1 \cdot\,\mathrm{d}\tau$. Because synchronization is slow, we keep the relative phases constant: $\delta_{ij} \approx$ const. Furthermore, we consider only the leading order interactions and set $\sigma_{ij} \approx \sigma(0) + 4\pi\tau$ in averaging the coupling. Then, the average radial dynamics become

$$\left\langle \frac{\mathrm{d}\bar{R}_i(\tau)}{\mathrm{d}\tau} \right\rangle = \frac{\langle 1 - \bar{R}_i(\tau)\rangle}{\tau_\kappa} + A\sin\left[\delta_{ij}(\tau) + \psi\right]. \tag{SI.31}$$

The hydrodynamic interaction causes a perturbation of the order of $A = O(a/d)$, which triggers a radial displacement $\langle 1 - \bar{R}_i\rangle = O(\tau_\kappa a/d)$. To the leading order in the hydrodynamic interaction the average dynamics of phases can be written as

$$\left\langle \frac{\mathrm{d}\phi_i(\tau)}{\mathrm{d}\tau} \right\rangle = 2\pi\left[1 + \langle 1 - \bar{R}_i(\tau)\rangle\right] + A\cos\left[\delta_{ij}(\tau) + \psi\right]. \tag{SI.32}$$

By exploiting $\tau_s \gg \tau_\kappa$, we obtain $\langle 1 - \bar{R}_i(\tau)\rangle = -\tau_\kappa A \sin(\delta_{ij} + \psi)$ from Eq. (SI.31), which yields the dynamics of phases on the synchronization timescale

$$\frac{\mathrm{d}\phi_i(\tau)}{\mathrm{d}\tau} = 2\pi + A\left[\cos\left(\phi_i(\tau) - \phi_j(\tau) + \psi\right) - 2\pi\tau_\kappa \sin\left(\phi_i(\tau) - \phi_j(\tau) + \psi\right)\right], \tag{SI.33}$$

where $A$ and $\psi$ are given by Eqs. (SI.24) and (SI.26), respectively, and we dropped the averaging indication.

#### B. Approximate calculation of synchronization rate and frequency gain

As shown in the main text, the synchronization rate $\tau_s^{-1}$ and frequency gain $\omega_s$ are given by

$$\tau_s^{-1} = 2A\left[\sin\psi + 2\pi\tau_\kappa\cos\psi\right], \tag{SI.34}$$

$$\omega_s = 2A\left[\cos\psi - 2\pi\tau_\kappa\sin\psi\right]. \tag{SI.35}$$

In the limit $\tau_\nu \ll 1$, we obtain $\tau_s^{-1}$ and $\omega_s$ by solving the history integrals [Eqs. (SI.18) and (SI.19)] approximately. To this end, we use integration by parts in Eq. (SI.18), yielding

$$\mathcal{X} = -\lim_{\tau\to\infty}\left[\mathcal{M}(\tau - \tau')\cos\{2\pi(\tau - \tau')\}\right]_0^\tau + \lim_{\tau\to\infty} 2\pi\int_0^\tau \mathcal{M}(\tau - \tau')\sin\left[2\pi(\tau - \tau')\right]\mathrm{d}\tau' \tag{SI.36}$$

with

$$\mathcal{M}(\tau) = \int\left(\bar{M}_1(\tau) - \frac{\bar{M}_3(\tau)}{2}\cos^2\beta\right)\mathrm{d}\tau. \tag{SI.37}$$

The memory kernels $\bar{M}_\alpha(\tau)$ [Eq. (SI.9)] peak when $\tau \approx \tau_\nu$ [see Fig. 3(a) of the main text]. For $\tau_\nu/\tau \ll 1$, the leading order term decays as $O[(\tau/\tau_\nu)^{-3/2}]$. Thus, $\mathcal{M}(\tau)$ decays to the leading order as $O[(\tau/\tau_\nu)^{-1/2}]$. Therefore, for short diffusion times ($\tau_\nu \ll 1$), the function $\mathcal{M}(\tau - \tau')$ decays rapidly, compared to the rotor motion, which is represented by $\cos\left[2\pi(\tau - \tau')\right]$. We use this to calculate the first term in Eq. (SI.36). We treat $\mathcal{M}(\tau - \tau')$ as changing instantaneously by letting $\tau' \to \tau$ in $\cos\left[2\pi(\tau - \tau')\right]$, but not in $\mathcal{M}(\tau - \tau')$. Thus,

$$-\lim_{\tau\to\infty}\left[\mathcal{M}(\tau - \tau')\cos\{2\pi(\tau - \tau')\}\right]_0^\tau \approx \lim_{\tau\to\infty}\left[\mathcal{M}(\tau) - \mathcal{M}(0)\right] = \frac{\tau_\nu}{32\pi}\left(5 + \cos 2\beta\right), \tag{SI.38}$$

where we calculated $\mathcal{M}(\tau)$ and its limit with *Mathematica*.

In the second term of Eq. (SI.36) the integral over $\mathcal{M}(\tau)$ is taken, which has an $O[(\tau/\tau_\nu)^{1/2}]$ contribution in the long time. Thus, assuming its rapid decay, compared to the rotor motion, is not justified. Instead, we follow Theers and Winkler [2] and develop the memory kernels $\bar{M}_1 = 1/12 \times [\tau_\nu/(\pi\tau)]^{3/2} + O[(\tau_\nu/\tau)^{5/2}]$ and $\bar{M}_3 = 0 + O[(\tau_\nu/\tau)^{5/2}]$ to the leading order in $\tau_\nu/\tau \ll 1$, which gives

$$\lim_{\tau\to\infty} 2\pi\int_0^\tau \mathcal{M}(\tau - \tau')\sin\left[2\pi(\tau - \tau')\right]\mathrm{d}\tau' \approx -\lim_{\tau\to\infty}\frac{\tau_\nu^{3/2}}{3\sqrt{\pi}}\int_0^\tau \frac{\sin\left[2\pi(\tau - \tau')\right]}{\sqrt{\tau - \tau'}}\mathrm{d}\tau' = -\frac{\tau_\nu^{3/2}}{6\sqrt{\pi}}, \tag{SI.39}$$

where we used the Fresnel $S$ integral. Using Eqs. (SI.38) and (SI.39) in Eq. (SI.36) gives

$$\mathcal{X} \approx \frac{\tau_\nu}{32\pi}\,(5+\cos 2\beta) - \frac{\tau_\nu^{3/2}}{6\sqrt{\pi}}. \tag{SI.40}$$

Repeating the calculations for the history integral (SI.19) yields

$$\mathcal{Y} \approx \frac{\tau_\nu^{3/2}}{6\sqrt{\pi}}. \tag{SI.41}$$

Using Eqs. (SI.40) and (SI.41) in Eqs. (SI.24) and (SI.26) we obtain

$$\tau_{\rm s}^{-1} \approx \frac{3\pi^2\gamma\tau_\kappa}{2}(5+\cos 2\beta) + 4\pi^{3/2}\gamma\sqrt{\tau_\nu}(1-2\pi\tau_\kappa) \tag{SI.42}$$

and

$$\omega_{\rm s} \approx \frac{3\pi\gamma}{4}(5+\cos 2\beta) - 4\pi^{3/2}\gamma\sqrt{\tau_\nu}(1+2\pi\tau_\kappa) \tag{SI.43}$$

from Eqs. (SI.34) and (SI.35), respectively. For $\beta = 0$, the limit $\tau_\nu \longrightarrow 0$ corresponds to the result of Ref. [3]. In the limit $\tau_\kappa \longrightarrow 0$, the correction terms proportional to $\sqrt{\tau_\nu}$ correspond to the result given in Ref. [2].

### IV. LINEAR STABILITY ANALYSIS OF METACHRONAL WAVES

Here we calculate the linear stability of metachronal waves, in periodic chains of $N$ phase oscillators with lattice constant $\ell$ (see Fig. 5 of the main text). To implement the periodic boundary conditions, we use a cutoff radius of half the system size, such that each oscillator interacts with the same number of neighbors in both directions of the chain. Metachronal waves have constant phase shifts $\phi_i - \phi_{i+1} = \ell k$ with wave number $k$ and $|\ell k| = 2\pi K/N$, where $K$ is an integer. We transform the phases as $\varphi_i(\tau) = \phi_i(\tau) - [2\pi + \Omega(k)]\tau$, where the frequency gain $\Omega$ of metachronal waves with wave number $k$ is given by

$$\Omega(k) = \sum_{j=1}^{m} \omega_{{\rm s},0j} \cos\left(j\ell k\right). \tag{SI.44}$$

Here $m = (N-1)/2$ for odd $N$, $m = N/2 - 1$ for even $N$, $\omega_{{\rm s},ij} = 2A_{ij}\left[\cos\psi_{ij} - 2\pi\tau_\kappa \sin\psi_{ij}\right]$, $A_{ij} = A[a/d_{ij}, (d_{ij})^2/(\nu T)]$, and $\psi_{ij} = \psi[(d_{ij})^2/(\nu T)]$, where $d_{ij} = \ell|i-j|$ with $i, j \in \{0, \ldots, N-1\}$ and $j \neq i$. The dynamic equations of the transformed phases are written as

$$\frac{{\rm d}\varphi_i}{{\rm d}\tau} = -\Omega(k) + \frac{1}{2}\sum_{j\neq i}\left[\omega_{{\rm s},ij}\cos(\varphi_i - \varphi_j) - \tau_{{\rm s},ij}^{-1}\sin(\varphi_i - \varphi_j)\right], \tag{SI.45}$$

where $\tau_{{\rm s},ij}^{-1} = 2A_{ij}\left[\sin\psi_{ij} + 2\pi\tau_\kappa\cos\psi_{ij}\right]$. In the periodic chain with phases $\varphi_i - \varphi_{i+1} = \ell k$, the sum in Eq. (SI.45) gives $\Omega$. Thus, metachronal waves are stationary solutions of Eq. (SI.45).

We obtain the dynamics of a small perturbation of metachronal waves $\boldsymbol{\Delta\varphi}(\tau) = [\Delta\varphi_0(\tau), \ldots, \Delta\varphi_{N-1}(\tau)]$, by linearization of Eq. (SI.45) as

$$\frac{\rm d}{{\rm d}\tau}(\boldsymbol{\Delta\varphi}) = \mathbf{J}(k)\cdot\boldsymbol{\Delta\varphi}, \tag{SI.46}$$

where the components of the Jacobi matrix $\mathbf{J}$ are given by

$$J_{ij}(k) = \frac{\partial}{\partial\varphi_j}\left(\frac{{\rm d}\varphi_i}{{\rm d}\tau}\right)\bigg|_{\varphi_j=-j\ell k}. \tag{SI.47}$$

Due to the periodic boundary conditions and the periodic metachronal waves, the Jacobi matrix $\mathbf{J}$ is a circulant matrix, for which the eigenvalues can be computed analytically [4]. For this, we only need the components

$$J_{0j} = \frac{\partial}{\partial\varphi_j}\left\{\frac{1}{2}\sum_{j\neq 0}\left[\omega_{{\rm s},0j}\cos(\varphi_0 - \varphi_j) - \tau_{{\rm s},0j}^{-1}\sin(\varphi_0 - \varphi_j)\right]\right\}_{\varphi_j=-j\ell k}. \tag{SI.48}$$

For the zeroth component, we have

$$J_{00} = -\frac{1}{2}\sum_{j\neq 0} \omega_{\mathrm{s},0j} \sin(\varphi_0 - \varphi_j)\Big|_{\varphi_j=-j\ell k} - \frac{1}{2}\sum_{j\neq 0} \tau^{-1}_{\mathrm{s},0j} \cos(\varphi_0 - \varphi_j)\Big|_{\varphi_j=-j\ell k}, \tag{SI.49}$$

The interaction ranges symmetrically in either direction of the chain. Thus,

$$\begin{aligned} J_{00} =& -\frac{1}{2}\sum_{j=1}^{m} \left[\omega_{\mathrm{s},0j} \sin(\varphi_0 - \varphi_j) + \omega_{\mathrm{s},0N-j} \sin(\varphi_0 - \varphi_{N-j})\right]_{\varphi_j=-j\ell k} \\ & -\frac{1}{2}\sum_{j=1}^{m} \left[\tau^{-1}_{\mathrm{s},0j} \cos(\varphi_0 - \varphi_j) + \tau^{-1}_{\mathrm{s},0N-j} \cos(\varphi_0 - \varphi_{N-j})\right]_{\varphi_j=-j\ell k}, \end{aligned} \tag{SI.50}$$

where $m = (N-1)/2$ for odd $N$ and $m = N/2 - 1$ for even $N$. Due to the periodic boundary conditions and the wave state, we can set $\omega_{\mathrm{s},0N-j} = \omega_{\mathrm{s},0j}$, $\tau^{-1}_{\mathrm{s},0N-j} = \tau^{-1}_{\mathrm{s},0j}$, $\phi_0 - \phi_j = j\ell k$, and $\phi_0 - \phi_{N-j} = -j\ell k$. Thus,

$$\begin{aligned} J_{00} &= -\frac{1}{2}\sum_{j=1}^{m} \left[\omega_{\mathrm{s},0j} \sin(j\ell k) + \omega_{\mathrm{s},0j} \sin(-j\ell k)\right] - \frac{1}{2}\sum_{j=1}^{m} \left[\tau^{-1}_{\mathrm{s},0j} \cos(j\ell k) + \tau^{-1}_{\mathrm{s},0j} \cos(-j\ell k)\right] \\ &= -\sum_{j=1}^{m} \tau^{-1}_{\mathrm{s},0j} \cos(j\ell k). \end{aligned} \tag{SI.51}$$

The $j \neq 0$ components of $J_{0j}$ can be written as

$$\begin{aligned} J_{0j} =& \frac{1}{2}\left[\omega_{\mathrm{s},0j} \sin(j\ell k) + \tau^{-1}_{\mathrm{s},0j} \cos(j\ell k)\right] \Theta(m-j) \\ &+ \frac{1}{2}\left\{\omega_{\mathrm{s},0N-j} \sin\left[-(N-j)\ell k\right] + \tau^{-1}_{\mathrm{s},0N-j} \cos\left[-(N-j)\ell k\right]\right\} \Theta[j-(N-m)], \end{aligned} \tag{SI.52}$$

where $\Theta(x)$ is the Heaviside function. The eigenvalues $\lambda_n$ of the circulant matrix $\mathbf{J}$ are given by

$$\lambda_n = \sum_{j=0}^{N-1} J_{0j} \exp\left(\mathrm{i}\frac{2\pi nj}{N}\right) = \sum_{j=0}^{N-1} J_{0j} \cos\frac{2\pi nj}{N} + \mathrm{i}\sum_{j=0}^{N-1} J_{0j} \sin\frac{2\pi nj}{N}, \tag{SI.53}$$

where $\mathrm{i} = \sqrt{-1}$ and $n \in \{0, \dots, N-1\}$. Using Eq. (SI.51) and Eq. (SI.52) in Eq. (SI.53), we obtain the real part as

$$\operatorname{Re}\lambda_n = -\sum_{j=1}^{m} \tau^{-1}_{\mathrm{s},0j} \cos(j\ell k) + \sum_{j=1}^{N-1} J_{0j} \cos\frac{2\pi nj}{N} \tag{SI.54}$$

$$= -\sum_{j=1}^{m} \tau^{-1}_{\mathrm{s},0j} \cos(j\ell k) + \frac{1}{2}\sum_{j=1}^{m} \left[\omega_{\mathrm{s},0j} \sin(j\ell k) + \tau^{-1}_{\mathrm{s},0j} \cos(j\ell k)\right] \cos\frac{2\pi nj}{N} \tag{SI.55}$$

$$+ \frac{1}{2}\sum_{j=1}^{m} \left[\omega_{\mathrm{s},0j} \sin(-j\ell k) + \tau^{-1}_{\mathrm{s},0j} \cos(-j\ell k)\right] \cos\frac{2\pi n(N-j)}{N}. \tag{SI.56}$$

Thus, by symmetry, the real part of the eigenvalues of the Jacobian $\mathbf{J}$ is given by

$$\operatorname{Re}\lambda_n(k) = -\sum_{j=1}^{m} \tau^{-1}_{\mathrm{s},0j} \cos(j\ell k) + \sum_{j=1}^{m} \tau^{-1}_{\mathrm{s},0j} \cos(j\ell k) \cos\frac{2\pi nj}{N}. \tag{SI.57}$$

### V. NUMERICAL METHODS

#### A. Numerical integration of the rotor dynamics

To simulate the rotor dynamics [Eq. (3) of the main text], we integrate the system of tangential [Eq. (SI.10)] and radial [Eq. (SI.11)] dynamics by the Euler method with numerical time step $\Delta\tau = 0.001$. In each time step, we evaluate the history

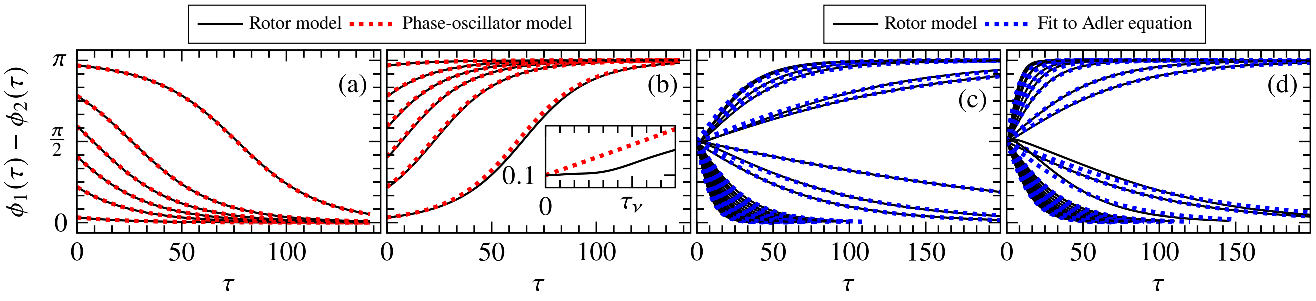


Fig. S1. (a)-(b) Time series of phase difference $\delta(\tau) = \phi_1(\tau) - \phi_2(\tau)$ for various initial conditions $\delta(0)$, according to the rotor model [Eqs. (SI.10) and (SI.11), black solid curves] and to the phase-oscillator model [Eqs. (SI.59) and (SI.34), red dotted curves], respectively. Data are generated with $\tau_\nu = 1$, $\tau_\kappa = 0.4$, $\gamma = 0.01$, $\bar{R}_1(0) = \bar{R}_2(0) = 1$, $\beta = 0$ (a), and $\beta = \pi/2$ (b). (c)-(d) Time series $\delta(\tau)$ corresponding to the data shown in Fig. 4(c) of the main text, with (c) the data for $\beta = 0$ and (d) the data for $\beta = \pi/2$, respectively. Here, black solid curves are data of the rotor model and blue dotted curves are the corresponding fit to the solution of the Adler equation (SI.59). The initial conditions for the rotor model are $\delta(0) = \pi/2$ and $\bar{R}_1(0) = \bar{R}_2(0) = 1$. For better visibility we only show parts of some of the time series.

functionals (SI.12) and (SI.13), respectively, using the trapezoidal rule with the history of phases $\phi_i(\tau')$ obtained by the Euler integration. The time step used determines the number of sampling points for the numerical integration of the history functionals. In our case, there are a thousand sampling points per intrinsic period. The memory kernels (SI.9) are peaked when $\tau \approx \tau_\nu$ and subsequently approach zero as $\bar{M}_1 = O[(\tau_\nu/\tau)^{3/2}]$ and $\bar{M}_3 = O[(\tau_\nu/\tau)^{5/2}]$, respectively. Thus, for small $\tau_\nu$ the temporal resolution might not be sufficient to capture the peak of the memory kernels accurately. Therefore, when running the simulation with $\tau_\nu \ll 1$, we split the integrals in two parts. One captures the long tail, the other the short peak.

The integrals in the history functionals (SI.12) and (SI.13) have the form $\int_0^\tau G(\tau-\tau')f[\phi(\tau')]\mathrm{d}\tau'$. For $\tau_\nu < 0.1$, we split the integrals according to

$$\int_0^\tau G(\tau-\tau')f[\phi(\tau')]\mathrm{d}\tau' = \int_0^{\tau-2\tau_\nu} G(\tau-\tau')f[\phi(\tau')]\mathrm{d}\tau' + \int_{\tau-2\tau_\nu}^{\tau} G(\tau-\tau')f[\phi(\tau')]\mathrm{d}\tau'. \tag{SI.58}$$

Here, the first term on the right captures the long tail, and the second term captures the short peak. For the long time integral we use the sample points directly obtained from the simulation. For the second integral, which contains the short peak, we refine the temporal resolution of $\tau'$ so that $2\times10^3$ samples are contained within the range $[\tau-2\tau_\nu, \tau]$. For $\tau_\nu \ll 1$, the phases $\phi_i(\tau')$ are approximately constant in the range $[\phi_i(\tau-2\tau_\nu), \phi_i(\tau)]$. By linear interpolation we increase the resolution of $\phi_i(\tau')$ to include $2\times10^3$ samples in this range. During the early stages of the simulation, when $\tau < 2\tau_\nu$, we do not split the integral and refine the data so that $[0,\tau]$ and $[\phi_i(0), \phi_i(\tau)]$ contain $2\times10^3$ samples, respectively.

#### B. Numerical integration of the history integrals

To calculate the amplitude $A$ [Eq. (SI.24)] and the phase delay $\psi$ [Eq. (SI.26)] of the phase-oscillator model, we solve the history integrals (SI.22) using the Simpson rule. To this end, we truncate the integration at the finite time $\tau = 10^4\tau_\nu$. At this time, the leading order term of the memory kernel has dropped to the order $O(10^{-6})$.

Figure S1(a)-(b) shows a comparison between the rotor model and the phase-oscillator model with the vorticity diffusion time equal to the period of rotor motion $\tau_\nu = 1$. The data shows that the phase difference in the rotor model (black solid curves) is captured to a good approximation by the solution of the Adler equation with identical frequencies (red dotted curves)

$$\delta(\tau) = 2\arctan\left[\tan\left(\frac{\delta(0)}{2}\right)\exp\left(-\frac{\tau}{\tau_\mathrm{s}}\right)\right], \tag{SI.59}$$

where $\tau_\mathrm{s}^{-1}$ is given by Eq. (SI.34).

#### C. Extraction of synchronization rate from rotor model data

In Fig. 4(c) of the main text we compare the synchronization rate according to the phase-oscillator model to the data of the full rotor model. To obtain the synchronization rate we extract time series of phase difference $\delta(\tau) = \phi_1(\tau) - \phi_2(\tau)$ from simulations of the rotor model with parameters as in Fig. 4(c) of the main text. The initial conditions of each simulation are $\delta(0) = \pi/2$

and $\bar{R}_1(0) = \bar{R}_2(0) = 1$. From the time series $\delta(\tau)$ we obtain the synchronization rate $\tau_s^{-1}$ by fit onto the solution of the Adler equation (SI.59). Figure S1(c)-(d) shows all time series and corresponding fit functions.

[1] M. R. Maxey and J. J. Riley, Equation of motion for a small rigid sphere in a nonuniform flow, The Physics of Fluids **26**, 883 (1983).
[2] M. Theers and R. G. Winkler, Synchronization of rigid microrotors by time-dependent hydrodynamic interactions, Physical Review E **88**, 023012 (2013).
[3] T. Niedermayer, B. Eckhardt, and P. Lenz, Synchronization, phase locking, and metachronal wave formation in ciliary chains, Chaos **18**, 1 (2008).
[4] M. Mehta, *Matrix Theory: Selected topics and useful results* (Les Éditions de physique, 1989).
